# A four-population model of *Diaphorina citri* and *Tamarixia radiata* with citrus flushing dynamics and impulsive parasitoid releases

**DOI:** 10.64898/2026.07.22.740061

**Authors:** Sebastián Rodríguez-Falcón, Heliana Arias-Castro, Javier Galeano, Luciano Stucchi

## Abstract

*Diaphorina citri* is the primary vector of Huanglongbing (HLB), a devastating disease that affects global citrus production. Effective biological control using the parasitoid *Tamarixia radiata* represents a sustainable alternative to chemical insecticides, but its efficacy depends heavily on environmental variables and resource availability. In this work, we develop a four-population mathematical model to analyze the dynamics of *D. citri* in the presence of *T. radiata*, a known parasitoid. Our model incorporates the cyclical and periodic behavior of citrus flushing (new shoots), providing a realistic representation of resource-limited dynamics as observed in field conditions. Through comparative simulations of three scenarios, without parasitoid, a single initial introduction, and periodic augmentative releases, we characterize and compare the impact of *T. radiata* as a biological control agent. Our results show that periodic releases maintain pest suppression, whereas a single introduction only delays pest recovery. By aligning theoretical modeling with ecological reality, this framework supports the role of *T. radiata* in pest suppression and provides a baseline for future work on optimal and cost-effective release strategies.

## 1. Introduction

Citrus is one of the most economically important fruit crops worldwide, and its production is severely threatened by Huanglongbing (HLB), or citrus greening disease, widely regarded as the most destructive disease affecting the crop today [1, 2]. The name derives from the Chinese *huanglongbing* (“yellow dragon disease”), in reference to the characteristic yellowing of infected shoots. HLB is a bacterial disease of citrus caused by a phloem-restricted, Gram-negative bacterium of the genus *Liberibacter*, which colonizes the sieve tubes of the plant phloem. Three species have been associated with the disease [1], of which *Candidatus* Liberibacter asiaticus (CLas) is the most widespread and the most damaging. The provisional status designation “*Candidatus*” reflects the fact that the bacterium has not been stably cultured *in vitro*, a limitation that has greatly hindered its study and the development of curative therapies.

The bacterium spreads through two main routes. The first, and epidemiologically most important, is transmission by the Asian citrus psyllid *Diaphorina citri* (Hemiptera: Liviidae), the primary vector of the pathogen associated with HLB. The insect acquires CLas while feeding on the phloem of infected trees and subsequently inoculates healthy plants during feeding [2, 3]. Transmission is persistent and propagative, meaning that the bacterium multiplies within the insect before being transmitted [4], and nymphs that acquire CLas generally become more efficient vectors than adults [5]. The second route is the grafting of infected plant material, which, together with the international trade of nursery plants and ornamental Rutaceae, drives the long-distance spread of the disease between regions. Native to the Asian subcontinent, *D. citri* has expanded its geographic range dramatically over the last decades, invading citrus-producing regions across Asia [6], the Americas [7], and more recently other continents [3]. Once established, *D. citri* populations can reach extremely high densities on citrus flush, and their close dependence on the emergence of new shoots has been repeatedly documented as a central driver of population growth [8, 9].

Because HLB has no effective cure once a tree becomes infected, management strategies rely almost entirely on controlling the psyllid vector, either chemically or biologically [10]. Chemical control, although widely used, is costly, environmentally disruptive, and increasingly compromised by insecticide resistance. Biological control constitutes a more sustainable alternative, and among the natural enemies of *D. citri*, the ectoparasitoid *Tamarixia radiata* (Hymenoptera: Eulophidae) stands out as the most effective and widely employed agent in classical and augmentative biological control programs worldwide [10]. Other parasitoid species, such as *Diaphorencyrtus aligarhensis* (Hymenoptera: Encyrtidae), have also been reported attacking *D. citri* nymphs, generally at much lower parasitization rates than *T. radiata* [11].

The effectiveness of *T. radiata* as a biological control agent depends strongly on the temporal availability of its host, which is in turn tightly coupled to the phenology of citrus flushing. Because *D. citri* nymphs develop exclusively on young shoots, the cyclical and often irregular production of new flush imposes a strongly periodic, resource-limited structure on the psyllid population, and consequently on any parasitoid that depends on it. Field studies in Puerto Rico have shown that both psyllid abundance and the incidence of parasitism by *T. radiata* track citrus flushing cycles closely, with the greatest and most prolonged parasitism occurring where new flush was most abundant and persistent [12]. Capturing this resource periodicity, at least in a qualitative sense, is therefore an important feature for any mathematical model that aims to describe pest dynamics realistically and to guide the design of effective parasitoid release strategies.

Population dynamics models offer a natural framework to formalize these interactions and to explore, in a quantitative way, the conditions under which biological control can suppress pest outbreaks. In particular, generalized logistic-mutualistic models, originally proposed to describe systems that combine several types of ecological interactions within a single mathematical structure [13], have proven useful for studying multispecies agroecosystems in which mutualism, antagonism, and resource limitation act simultaneously. Stucchi et al. [14] extended this framework to a nursery-pollination system involving Caryophyllaceous plants, *Hadena* moths, and their parasitoid wasps, showing that the introduction of a parasitoid population can substantially stabilize an otherwise oscillatory or extinction-prone system, and that the net effect of a parasitoid on the whole community depends critically on the intensity of the predation term linking it to its host.

A complementary line of research has approached parasitoid-mediated biological control from the perspective of impulsive dynamical systems, in which continuous host-parasitoid interactions are punctuated by discrete, periodic releases of the natural enemy. This approach has been applied successfully to other agricultural pest systems with pronounced seasonality, such as the sugarcane borer *Diatraea saccharalis* and its egg and larval parasitoids, where periodic impulsive releases were shown to be an effective and practical alternative to continuous control strategies [15]. Field evidence from citrus agroecosystems similarly indicates that the impact of *T. radiata* on *D. citri* populations is highly variable and density-dependent, being modulated by abiotic mortality factors, climate, and interactions with other organisms such as ants [16, 17], which underscores the need for mechanistic models capable of capturing this variability rather than treating parasitoid efficacy as constant.

Building on this body of work, in the present paper we adapt the generalized model of García-Algarra et al. [13] and Stucchi et al. [14] to describe the interaction between four coupled populations in a citrus agroecosystem: the pest *D. citri*, citrus shoots, the parasitoid *T. radiata*, and a fourth compartment representing external agronomic or environmental stimuli (e.g., temperature, irrigation, or other factors) that modulate shoot production. The nonlinear coupling between the shoot and stimuli compartments generates sustained, resource-limited fluctuations that qualitatively reproduce the boom-and-bust pattern of citrus flushing documented under field conditions [12, 8], while the antagonistic term between pest and parasitoid captures the regulatory role that *T. radiata* is known to exert on *D. citri* populations [10, 11]. In addition, following the impulsive control framework developed for other host-parasitoid agroecosystems [15], we introduce impulsive perturbations in the parasitoid population to represent periodic augmentative releases, a strategy increasingly used in operational biological control programs [16, 17]. By comparing the system’s behavior in the absence of the parasitoid, under a single initial introduction, and under sustained periodic releases, we aim to characterize and compare the impact of *T. radiata* as a biological control agent and to provide a baseline framework for the future design of optimal, cost-effective release schedules that minimize *D. citri* infestations under realistic field conditions.

The remainder of this paper is organized as follows. Section 2 introduces the generalized four-population model, details the ecological justification for each interaction term, and formulates the impulsive release scheme. Section 3 presents the resulting population dynamics under the three release scenarios described above. Section 4 discusses the implications of these results for biological control practice and outlines directions for future work on optimal release strategies.

## 2. Methods

Following the framework introduced by Stucchi et al. [14], we consider a system of nonlinear ordinary differential equations describing the interaction between four populations in a citrus agroecosystem: the pest *Diaphorina citri*, plant shoots, the parasitoid *Tamarixia radiata*, and external stimuli.

Let *x*_*i*_(*t*) denote the density of population *i* at time *t*. The general structure of the model is given by

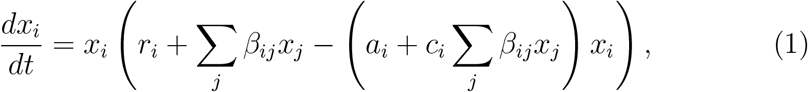

where *r*_*i*_ represents intrinsic growth rates, *a*_*i*_ self-limitation coefficients, *c*_*i*_ interaction-dependent regulation, and *β*_*ij*_ encodes the effect of population *j* on population *i*.

Following the approach of Stucchi *et al*., we construct the model by first identifying the main ecological interactions and then translating them into mathematical terms. In this framework, ecological interactions are encoded through the coefficients *β*_*ij*_, whose sign determines the nature of the interaction: positive values correspond to beneficial effects (e.g., resource availability), while negative values represent antagonistic interactions such as predation or competition. This allows the model to capture multiple ecological mechanisms within a unified mathematical structure, as in [14].

We consider four interacting populations: *Diaphorina citri* (*x*_1_), plant shoots (*x*_2_), the parasitoid *Tamarixia radiata* (*x*_3_), and external stimuli (*x*_4_). The system is simplified by retaining only the dominant pairwise interactions.

### Diaphorina citri and plant shoots

The pest population depends on the availability of plant shoots as a resource. This interaction contributes positively to the growth of *x*_1_ through the term *β*_12_*x*_2_, representing resource-driven reproduction. Conversely, the feeding activity of the pest negatively impacts the plant population, which is captured by the term *β*_21_*x*_1_ in the equation for *x*_2_.

### Diaphorina citri and parasitoids

The parasitoid *Tamarixia radiata* acts as a natural enemy of the pest. This interaction is antagonistic: parasitoids reduce the pest population, represented by a negative contribution *β*_13_*x*_3_ in the equation for *x*_1_. At the same time, parasitoids depend on the pest for reproduction, leading to a positive term *β*_31_*x*_1_ in the equation for *x*_3_.

### Plant shoots and external stimuli

External stimuli (e.g., environmental or agronomic factors) promote plant growth. This effect is modeled as a positive interaction *β*_24_*x*_4_ in the equation for *x*_2_. In contrast, plant abundance may reduce the relative impact of stimuli, leading to a negative feedback represented by *β*_42_*x*_2_ in the equation for *x*_4_.

### Self-limitation and saturation effects

Each population is subject to intraspecific competition and saturation effects. These are modeled by two parameters: *a*_*i*_, representing classical self-limitation, and *c*_*i*_, which accounts for the reduction of interaction efficiency at high population densities. Together, they produce the nonlinear term

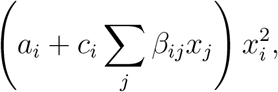

which limits unbounded growth.

In our case, the system is specified as follows:

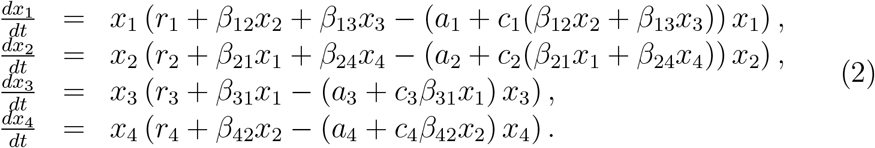

Here, *x*_1_, *x*_2_, *x*_3_, and *x*_4_ correspond to *Diaphorina citri*, plant shoots, *Tamarixia radiata*, and external stimuli, respectively.

To model periodic augmentative biological control, we introduce impulsive perturbations in the parasitoid population. Specifically, at discrete times *t* = *nT* , with *n* ∈ ℕ, the parasitoid density undergoes an instantaneous increase of magnitude Δ:

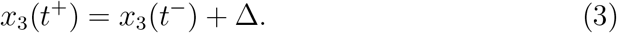

Between impulsive events, the system evolves continuously according to Eqs. (2).

The dynamics of the parasitoid population can be equivalently expressed as an impulsive differential equation:

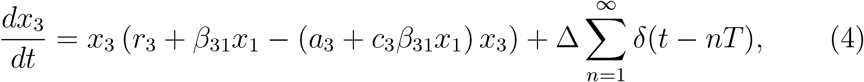

where *δ*(·) denotes the Dirac delta distribution.

This formulation makes explicit that the continuous dynamics is supplemented by discrete inputs occurring at regular time intervals. The numerical values we used in our simulations are shown in Table 1. Our code is published in [18].

**Table 1:** Model parameters and numerical values.

| Parameter | Numerical Value | Units |
| --- | --- | --- |
| $r_1$ | -0.20 | time<sup>-1</sup> |
| $r_2$ | -0.72 | |
| $r_3$ | -0.30 | |
| $r_4$ | 0.72 | |
| $b_{12}$ | 0.040 | population<sup>-1</sup> time<sup>-1</sup> |
| $b_{13}$ | -0.010 | |
| $b_{21}$ | -0.010 | |
| $b_{24}$ | 0.054 | |
| $b_{31}$ | 0.010 | |
| $b_{42}$ | -0.072 | |
| $a_1$ | $5.55 \times 10^{-4}$ | |
| $a_2$ | | |
| $a_3$ | | |
| $a_4$ | | |
| $c_1$ | $5.0 \times 10^{-3}$ | 1 |
| $c_2$ | | |
| $c_3$ | | |
| $c_4$ | | |
| $x_1(0)$ | 50 | population |
| $x_2(0)$ | 10 | |
| $x_3(0)$ | 15 | |
| $t_f$ | 100 | time |

## 3. Results

We analyze the population dynamics of the system in three stages. First, we consider a baseline scenario with no introduction of parasitoids and no subsequent releases. Then, we consider a scenario that incorporates a single initial population of parasitoids with no further releases. Our final scenario starts with an initial population of parasitoids and periodic releases to evaluate their impact on the system dynamics.

We show in Fig 1 the time evolution of the populations when no parasitoids are present. We see that the population of Diaphorina grows and oscillates following the same frequency established by the plant shoots and the stimuli dynamics.

**Figure 1:**
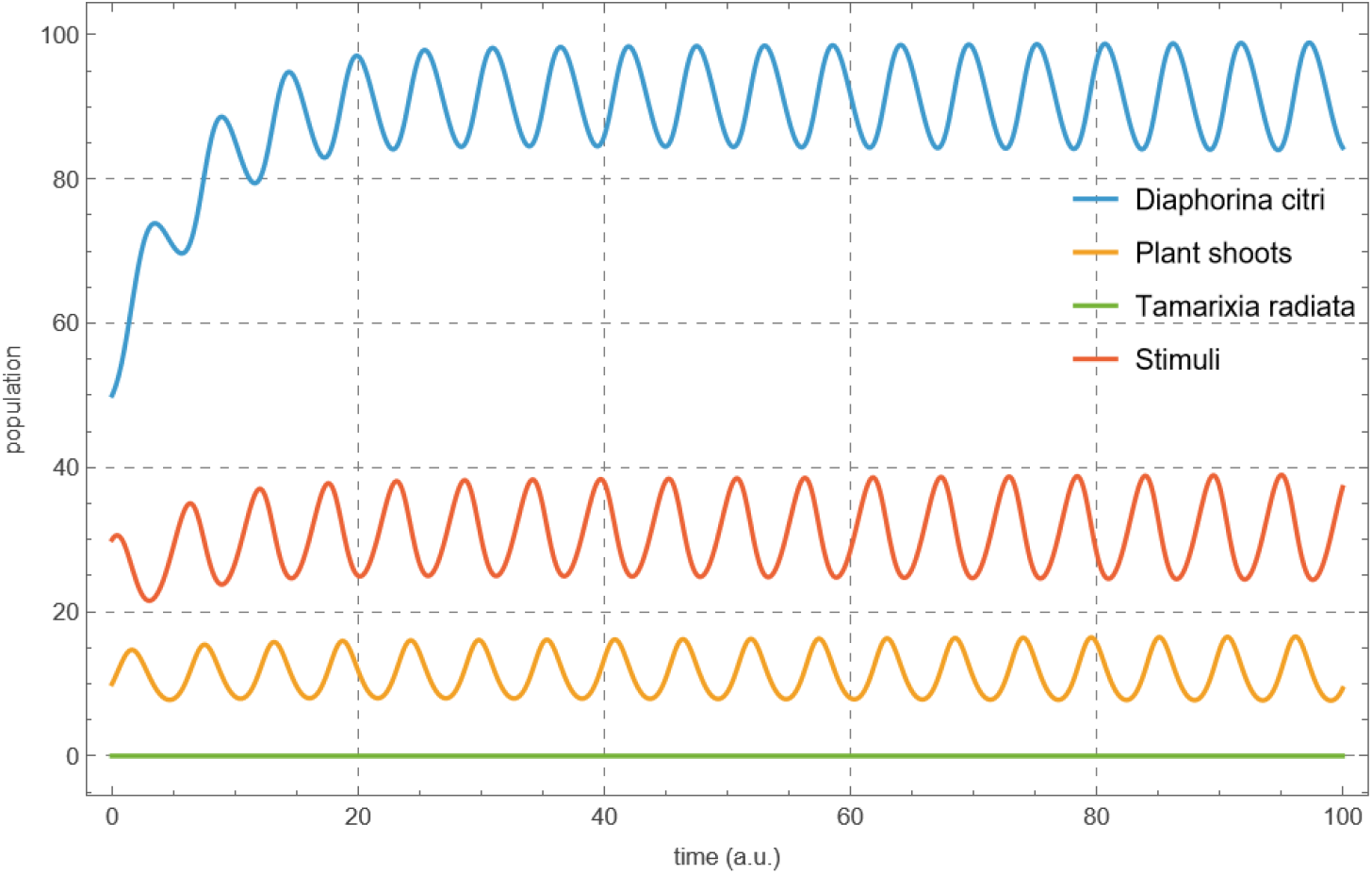
Time evolution of the four populations without parasitoid releases. No introduction of parasitoids leads to stable equilibrium characterized by high parasite densities. Without top-down regulation (i.e. with Δ = 35), the *Diaphorina c*. population rapidly reaches its carrying capacity and remains persistently elevated.

Figure 2 shows the time evolution of the system when an initial population of parasitoids is introduced, but no further releases are applied. In this case, the system evolves purely under its intrinsic nonlinear dynamics.

**Figure 2:**
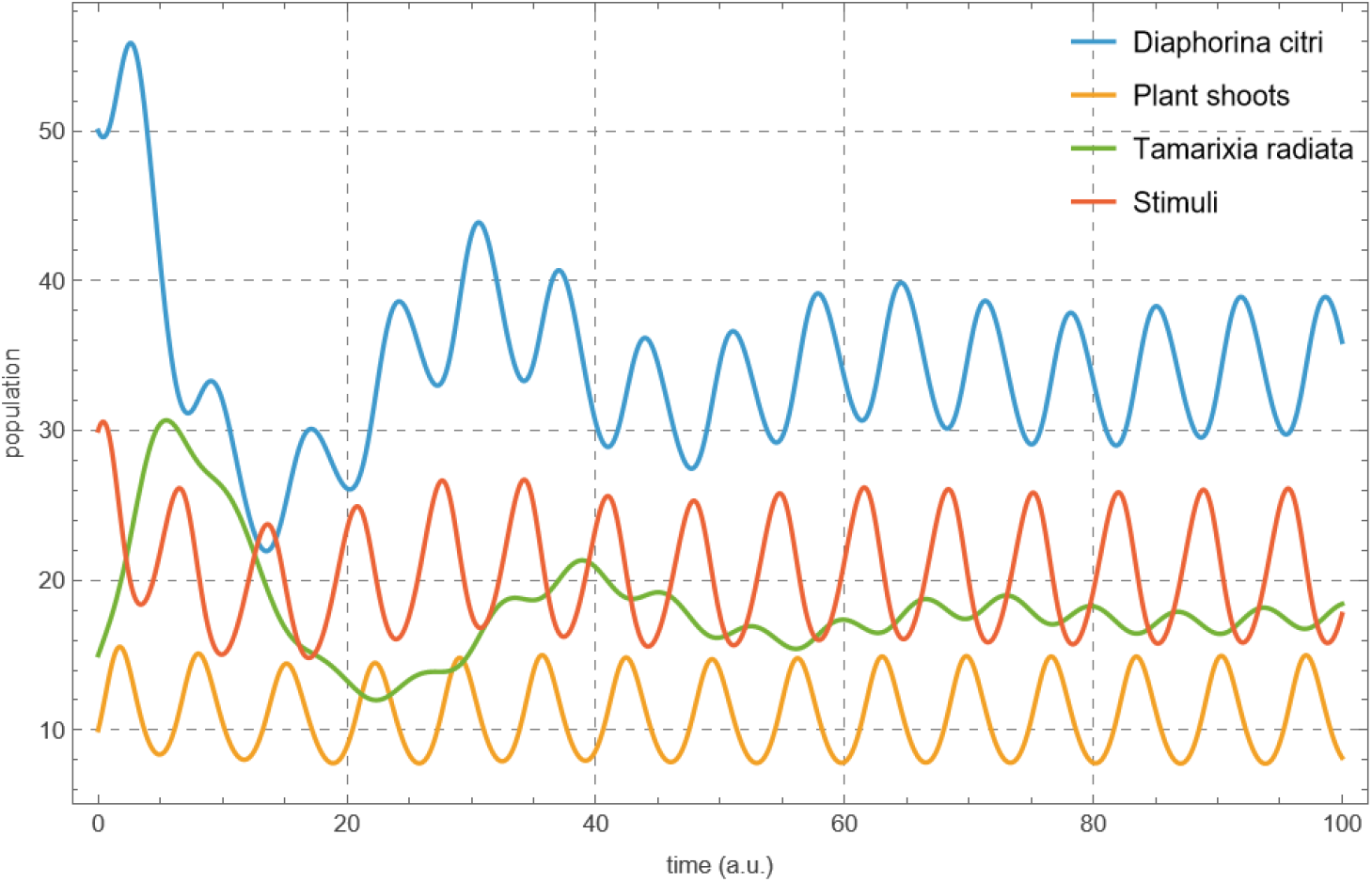
Time evolution of the four populations without periodic parasitoid releases. An initial introduction of parasitoids leads to transient suppression of the pest population, followed by recovery due to the decay of the parasitoid density.

The parasitoid population *x*_3_ initially increases due to the availability of the pest population *x*_1_, but subsequently declines as a result of self-limitation and reduced host availability. This leads to a transient suppression of the pest population, followed by a recovery phase once the parasitoid density decreases.

As a consequence, the system exhibits oscillatory behavior driven by the antagonistic interaction between the pest and the parasitoid. The plant population responds indirectly to these oscillations, reflecting the coupling between resource availability and pest pressure. The external stimulus variable evolves more smoothly and acts as a slower modulating component.

We now introduce periodic augmentative releases of parasitoids, modeled as impulsive increases of magnitude Δ = 35 applied at times *t* = *nT* , with *T* = 15.

Figure 3 shows the resulting dynamics. In contrast to the baseline scenario, the parasitoid population exhibits discontinuous jumps at each release time, preventing its decay to low levels.

**Figure 3:**
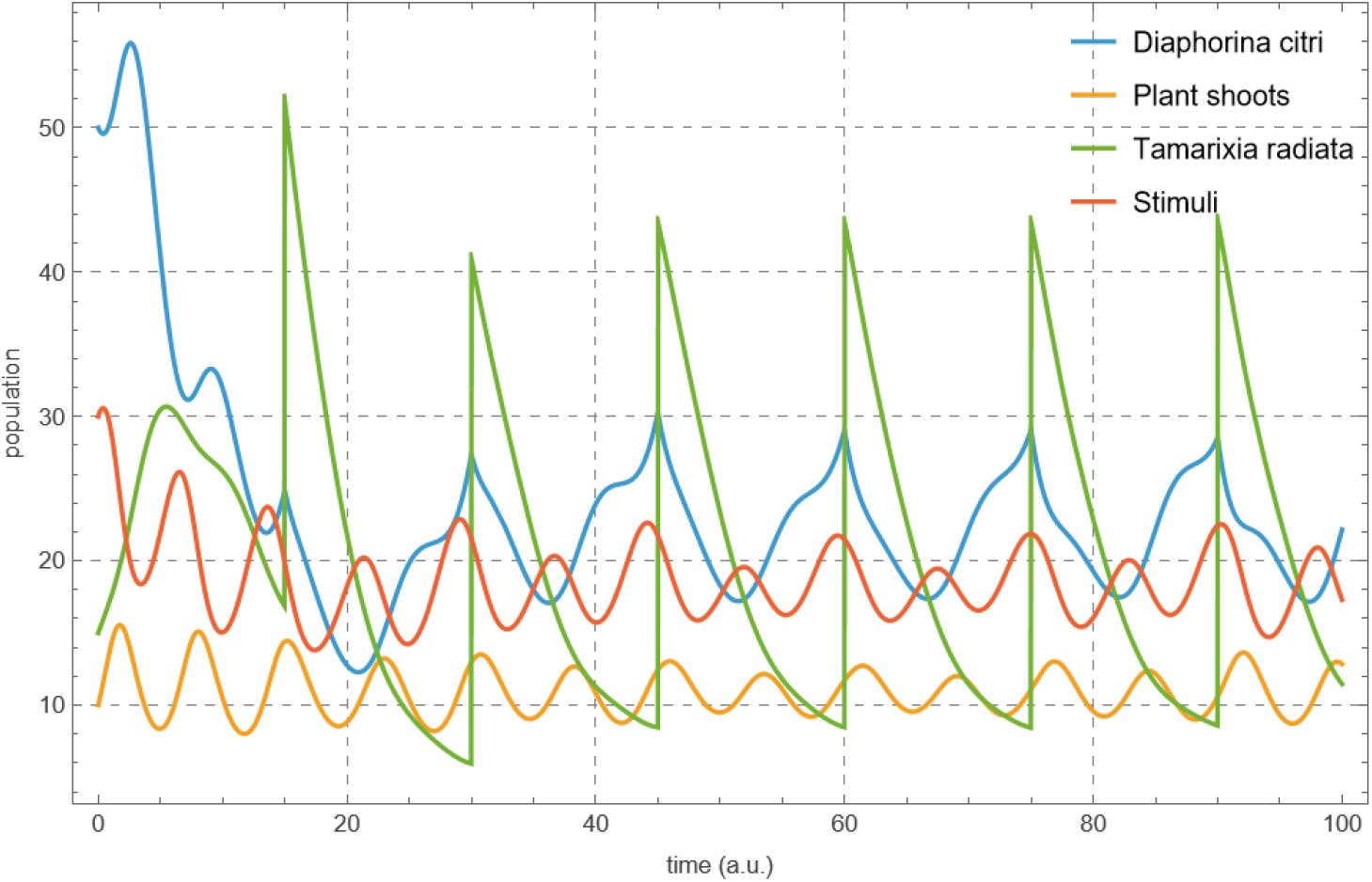
Time evolution of the system under periodic parasitoid releases with parameters (*T*, Δ). The parasitoid population exhibits discontinuous jumps at release times, sustaining its density and enabling long-term suppression of the pest population.

These repeated inputs sustain a higher average parasitoid density, leading to more persistent suppression of the pest population. Each pulse produces an immediate increase in *x*_3_, followed by a delayed decrease in *x*_1_, consistent with the predator–prey interaction structure of the system.

The plant population benefits indirectly from this control mechanism, as reduced pest pressure leads to higher average values of *x*_2_. At the same time, the system displays hybrid dynamics, combining continuous evolution with discrete perturbations.

Comparing both scenarios, we observe that periodic releases fundamentally alter the long-term behavior of the system. While the baseline dynamics are characterized by transient control followed by pest recovery, the introduction of periodic pulses maintains sustained regulation of the pest population.

The effectiveness of this control mechanism depends on the choice of the parameters (*T*, Δ). In particular, sufficiently frequent and/or strong releases can prevent large pest outbreaks, whereas weaker interventions may only delay them.

## 4. Conclusions

In this work, we adapted the generalized logistic-mutualistic framework of [14, 13] to describe the population dynamics of *Diaphorina citri* in the presence of its parasitoid, *Tamarixia radiata*, within a four-population citrus agroecosystem that also includes plant shoots and external environmental stimuli. Rather than imposing an explicit periodic forcing, the nonlinear coupling between the shoot and stimuli compartments generates sustained, resource-limited fluctuations that qualitatively reproduce the boom-and-bust pattern of citrus flushing reported under field conditions [12, 8], providing a more ecologically grounded representation of the resource constraints that shape psyllid population growth than a purely autonomous pest-parasitoid system would allow.

The comparative analysis of the three release scenarios considered here, namely the absence of the parasitoid, a single initial introduction, and sustained periodic augmentative releases, allowed us to characterize qualitatively different long-term regimes. In the absence of top-down regulation, the pest population stabilizes at persistently high densities. A single introduction of parasitoids produces only a transient suppression, followed by pest recovery as the parasitoid population decays due to self-limitation and reduced host availability. In contrast, periodic impulsive releases sustain a higher average parasitoid density and maintain long-term regulation of the pest population, consistent with the predator-prey structure underlying the antagonistic interaction between *D. citri* and *T. radiata*. These results support the general expectation that augmentative releases, rather than isolated introductions, are necessary to achieve durable suppression of *D. citri* under resource-limited, seasonally fluctuating conditions.

It is important to situate these theoretical results within the context of field evidence on the efficacy of *T. radiata*. Empirical studies have repeatedly shown that the impact of this parasitoid on *D. citri* populations is highly variable across sites and seasons, and is strongly modulated by abiotic mortality, climate, and biotic interactions such as ant-mediated disruption of natural enemies [16, 17]. The present model, in its current form, does not capture this variability explicitly, since the coefficients governing the pest-parasitoid interaction are held constant rather than being allowed to depend on environmental or biotic covariates. Bridging this gap between the qualitative regularities produced by the model and the quantitative variability observed in the field is a natural direction for future refinement.

Several other limitations of the present framework should be acknowledged. First, the parameter values used in the simulations were chosen to illustrate the qualitative behavior of the system rather than fitted to biological data specific to the *D. citri* –*T. radiata* system; incorporating parameter estimates derived from the literature, in the spirit of the empirically grounded calibration performed by [14], would strengthen the quantitative relevance of the model’s predictions. Second, the model treats each population as a single, undifferentiated compartment, without the stage structure (egg, nymph, adult) that characterizes the biology of both *D. citri* and *T. radiata* [3, 9]; a stage-structured extension could resolve delays between parasitism and host suppression that are not captured here. Third, the current formulation is deterministic, and does not account for demographic or environmental stochasticity, which are known to be relevant at the population densities and spatial scales typical of citrus orchards. Finally, although the impulsive release scheme introduced here provides a natural entry point for cost-based optimization, no explicit economic or operational cost function was formulated or optimized in this study.

Regarding future lines of research, the most direct extension of this work is the systematic exploration of the impulsive release parameters, the period *T* and the release magnitude Δ, following the impulsive and Floquet-theoretic approaches that have proven effective in other seasonal host-parasitoid agroecosystems [15]. Mapping the region of the (*T*, Δ) parameter space that guarantees sustained pest suppression, and identifying the combinations that minimize parasitoid input for a given suppression target, would provide the quantitative basis needed to design release schedules that are both biologically effective and economically efficient. Coupling such an analysis with field-calibrated parameters and an explicit representation of environmental variability would allow this framework to move from a qualitative proof of concept toward a practical decision-support tool for the management of *D. citri* in citrus-producing regions affected by HLB.

## 5. Authors contribution

Sebastián Rodríguez-Falcón: Software and writing - original draft (equal). Heliana Arias-Castro: Writing - review and editing (equal); Javier Galeano: Conceptualization and writing - review and editing (equal); Luciano Stucchi: Conceptualization and writing - review and editing (lead).

## 7. Acknowledgments

L.S. acknowledges financial support from the Centro de Investigación de la Universidad del Pacífico (CIUP). JG acknowledges financial support from the Spanish Ministry of Science, Innovation and Universities through the State Research Agency (Agencia Estatal de Investigación, AEI), under project NetLIFE-CODES (Grant PID2024-157869NB-I00).

